# Sequence-based TCR-Peptide Representations Using Cross-Epitope Contrastive Fine-tuning of Protein Language Models

**DOI:** 10.1101/2024.10.25.619698

**Authors:** Chiho Im, Ryan Zhao, Scott D. Boyd, Anshul Kundaje

## Abstract

Understanding T-Cell receptor (TCR) and epitope interactions is critical for advancing our knowledge of the human immune system. Traditional approaches that use sequence similarity or structure data often struggle to scale and generalize across diverse TCR/epitope interactions. To address these limitations, we introduce ImmuneCLIP, a contrastive fine-tuning method that leverages pre-trained protein language models to align TCR and epitope embeddings in a shared latent space. ImmuneCLIP is evaluated on epitope ranking and binding prediction tasks, where it consistently outperforms sequence-similarity based methods and existing deep learning models. Furthermore, ImmuneCLIP shows strong generalization capabilities even with limited training data, highlighting its potential for studying diverse immune interactions and uncovering patterns that improve our understanding of human immune recognition systems.

## 1 Introduction

T lymphocytes (T-Cells) are a key components of the adaptive immune response, surveying cells for the presence of foreign peptides within and responding to foreign epitopes displayed by antigen-presenting cells. This process is mediated by specific interactions between T-cell receptors (TCRs) and foreign peptides presented via the major histocompatibility complexes (pMHCs) [1]. Each TCR is a hetero-dimer composed of an *α* and *β* TCR chain, with three complementarity determining regions (CDR 1, 2, and 3) per chain. These CDRs have hypervariable sequences and determine the antigen specificity of the TCR. CDR1 and CDR2 interact primarily with the MHC, while CDR3, which interacts mostly with the peptide epitopes, dictates antigen specificity [2, 3]. Accurately predicting TCR-pMHC binding has significant implications for immunotherapy and understanding immune responses but remains challenging due to the high variability in TCR and epitope sequences.

Machine learning has advanced TCR-pMHC interaction prediction, using models based on Gaussian processes [4], random forests [5], nearest-neighbors using various distance metrics [6], and deep learning [7, 8, 9, 10, 11]. Early deep learning approaches incorporated biological priors into protein sequences using Atchley factors [12] and employed convolutional, long short-term memory (LSTM) networks (e.g. pMTnet) or transformer architectures (e.g. Pan-Pep). Models that incorporate TCR and pMHC structure have further improved predictions [13, 14], but the scarcity of reliable structural data has limited the broad applicability of these approaches. More recently, token-based language modeling methods using sequence alone have also shown great promise. For example, STAPLER employs masked language model pre-training on randomly paired TCR *α, β* CDR3 regions and epitope sequences before fine-tuning on full-length TCRs and their known binders [10]. TULIP constructs separate encoder-decoder blocks for the *α* CDR3, *β* CDR3, and epitope sequences and trains each decoder to predict its respective component given encodings from the other two blocks [11]. While these methods have achieved the state-of-the-art in predictive power, the paucity of TCR/epitope binding data is still a significant bottleneck and room for improvement remains.

Representations learned by self-supervised protein language models (PLMs) pre-trained on vast compendia of protein sequences have proven valuable for several downstream prediction tasks, especially in settings with limited labeled data. For example, embeddings from the ESM-2 PLM have resulted in improvements in protein structure prediction and coding variant effect prediction [15]. PLMs focused on antibody sequences, such as IgLM and AntiBERTa, as well as TCR sequences, such as TCR-BERT and TCRLang-Paired, have demonstrated their ability to encode features relevant for predicting immune interactions [16, 17, 18, 19]. Fine-tuning these PLMs on smaller labeled datasets for specific tasks has become a widely adopted effective strategy [20]. Notably, multi-modal contrastive learning methods such as Contrastive Language-Image Pre-training (CLIP), originally developed for aligning images with natural language text descriptions [21], have been successfully adapted for several multi-modal molecular interaction prediction tasks, as demonstrated by CLIPZyme for mapping enzymes to reactions and EAGLE for antibody-antigen interaction prediction [22, 23].

In this work, we introduce ImmuneCLIP, a scalable approach for contrastively fine-tuning PLMs to predict TCR/epitope binding. Our method aligns TCR and peptide embeddings in a shared latent space, capturing the diverse interactions within the human immune system. ImmuneCLIP outperforms sequence distance-based methods like tcrdist3 and surpasses current TCR-pMHC interaction prediction models. This sequence-based approach offers a powerful solution for multi-epitope binding prediction, which could in the future facilitate advancements in immunotherapy, vaccine design, and personalized medicine.

## 2 Methods

In this section, we outline the design and implementation of ImmuneCLIP, including how the model processes data for training and evaluation, generates learned representations of epitopes and T-cell receptors (TCRs), and employs contrastive learning to fine-tune embeddings. Our approach builds upon existing protein language models and applies parameter-efficient techniques to train the model while ensuring scalability and performance. Figure 1 illustrates an overview of the model architecture.

**Figure 1:**
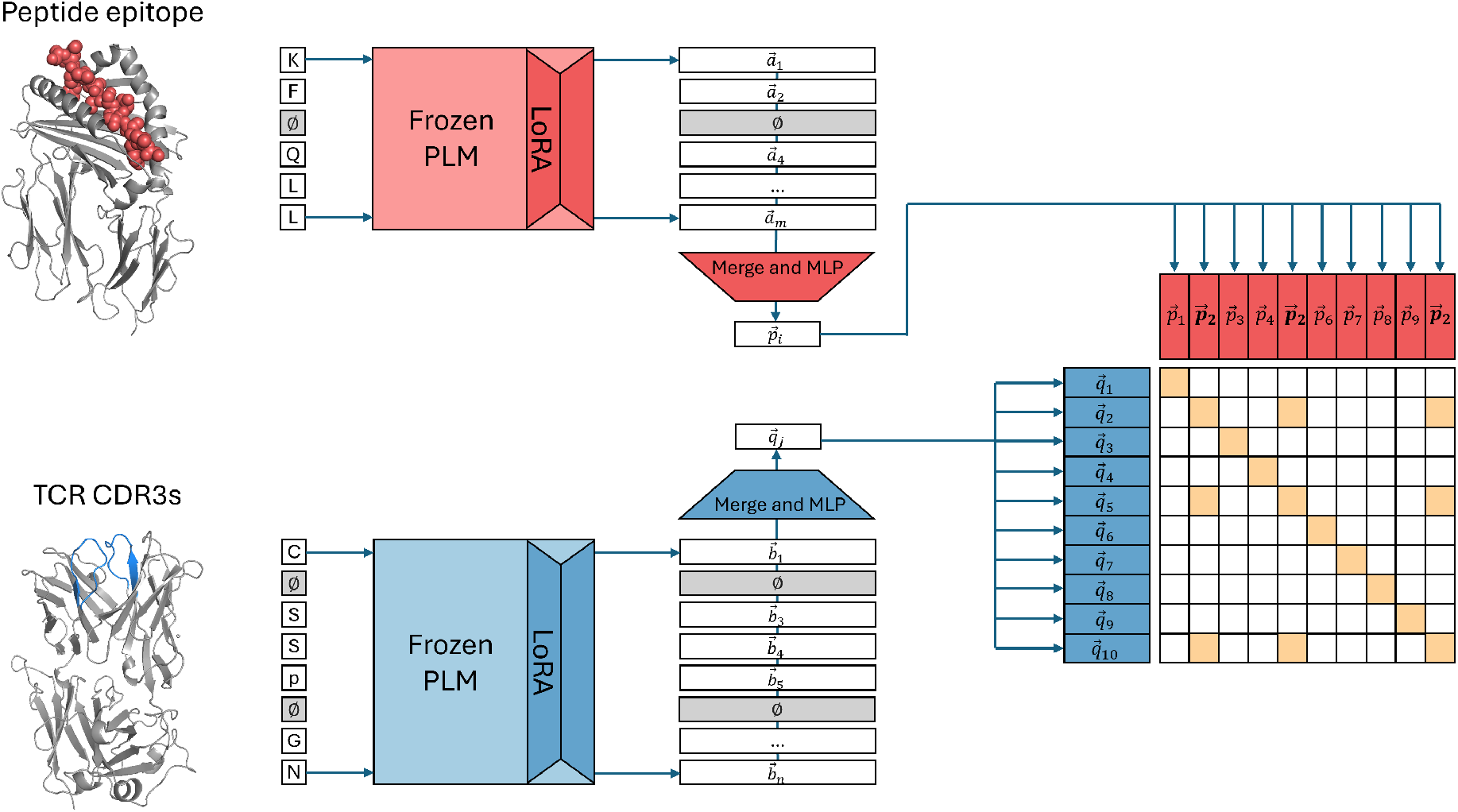
Overview of ImmuneCLIP architecture. Epitopes and T-Cell receptors (TCRs) are embedded separately using pre-trained protein language models (e.g., ESM-2 for epitopes and TCR-BERT for TCR CDR3 *α, β* chains), with parameter-efficient fine-tuning using LoRA. Embeddings from the final hidden layer are pooled and projected into a shared latent space using multi-layer perceptron (MLP) layers. The model is trained with a contrastive loss to maximize cosine similarity between embeddings of known binding pairs.

### 2.1 Dataset and Preprocessing

A number of curated TCR/epitope datasets exist, each offering an opinionated subset of the publicly available TCR and epitope sequencing space [7, 24, 25]. To train ImmuneCLIP, we selected the MixTCRPred dataset curated by Croce at al., which draws from IEDB, VDJdb, McPAS, and 10x Genomics [26, 27, 28, 29, 30]. The dataset matches epitope fragments to the paired *α* and *β* chain CDR3 sequences of the TCRs that binds them. Literature has shown that the CDR3 region is the most critical for epitope binding, with both the *α* and *β* CDR3s playing a role in forming the TCR binding site, making the MixTCRPred dataset an ideal choice.

The initial dataset contained 17,715 different pairs of TCR *α, β* CDR3 sequences and an epitope they bind to, consisting of 14,144 unique *α, β* CDR3 pairs and 146 unique epitopes, from both human and mouse origins. Filtering for duplicate (CDR3 *α*, CDR3 *β*, epitope) entries reduced the dataset size to 14,245 TCR/epitope pairs. Additionally, for our experiments, we focused exclusively on human TCR/epitope pairs, as the available mouse data was heavily skewed [Figure 2]. The final dataset consisted of 8439 unique human TCR/epitope pairs, which was then split into training, validation, and test sets with a ratio of 70:15:15. To further reduce the effects of skew in epitope counts and improve model performance on low-N epitopes during training, TCR/epitope pairs were upsampled during training as follows: For each epitope *e*_*i*_ of *p* unique epitopes, let 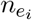 be the number of examples (receptors) for that epitope. The upsampled number 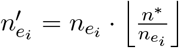, where 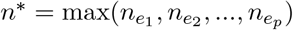.

**Figure 2:**
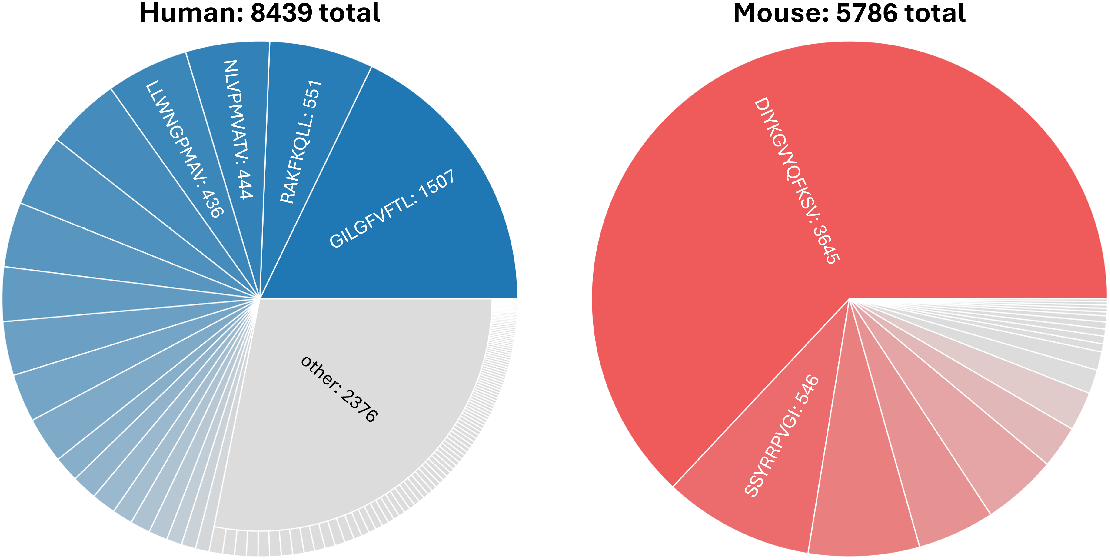
Breakdown of MixTCRPred dataset after duplicate filtering, separated by species of origin and sorted by the number of associated TCRs. The human set consists of 127 unique epitopes, with the top 20 epitopes illustrated and remainder grouped into “other”. The mouse set consists of 19 unique epitopes.

### 2.2 Generating Learned Representations using Protein Language Models

ImmuneCLIP generates separate learned representations for epitopes and TCRs. Epitopes are embedded using a pre-trained large protein language model (e.g. ESM-2 or ESM-3) [15, 31]. The CDR3 regions of TCR *α, β* chains are similarly embedded using a general protein language model or a TCR-specific language model (e.g. TCR-BERT or TCRLang) [18, 19]. We employ parameter-efficient fine-tuning using low rank adaptation (LoRA) to train ImmuneCLIP for TCR/epitope alignment. The pre-trained weights are frozen, and trainable low-rank matrices are inserted into the transformer layers, significantly reducing the number of trainable parameters while maintaining the capacity to learn task-specific features.

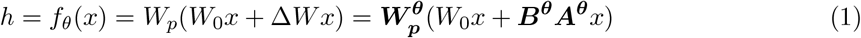

Formally, ImmuneCLIP creates an epitope embedding in 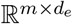, where *m* is the length of the epitope and *d*_*e*_ is the dimension of the language model used to embed epitopes. Similarly, the TCR embedding is in 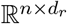, where *n* is the length of the TCR *α, β* chain CDR3, and *d*_*r*_ is the dimension of the language model used to embed T-cell receptors. Epitope and receptor sequences are tokenized into amino acids and fed into their respective LoRA-adapted protein language models (Equation 1), where *W*_0_ is the frozen PLM and matrices *A*^*θ*^ and *B*^*θ*^ of rank *r* are the LoRA matrices [32]. Following Sledzieski et al.’s approach for LoRA training of protein language models, adapters of rank 8 were added to the *key* and *value* weight matrices of the PLM self-attention layers [33]. The per-residue embeddings for epitopes and TCRs from the final hidden layer of the LoRA-adapted protein language models are then pooled across the residue dimension to create a fixed length vector for each embedding. The vectors are then projected onto the same lower-dimensional latent space (in 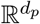, where *d*_*p*_ < max (*d*_*e*_, *d*_*r*_)) using one or more multi-layer perceptron (MLP) layers, 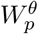. This facilitates the alignment of epitope and TCR sequence embeddings by enabling the identification of binding pairs through similarity measurements in the projected latent space.

During training, we partially mask out the epitope and receptor sequences to reduce the model overfitting [34, 35]. Many biological language models use *span masking* on their input sequences during pre-training and fine-tuning, where a random, continuous sub-string of the overall sequence is masked out during each iteration (usually ∼ 15% of the total sequence length) [15, 36]. Given that epitope and TCR CDR3 sequences are shorter than full-length protein sequences, we decided to mask each token individually, where each token is masked with probability *p* = 0.15, while ensuring that not every residue is masked. Masked residue embeddings are omitted from the pooling stage that is applied to the protein language model outputs.

### 2.3 Contrastive Learning for Epitope/TCR Representation Fine-Tuning

The protein language model representations of antigen epitopes and T-cell receptors in ImmuneCLIP are fine-tuned using a CLIP-style contrastive learning architecture [21, 37]. Let the representation of an epitope *E* be denoted as 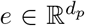 and the representation of a T-cell receptor *R* as 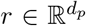. Our objective tries to learn a function *g*(*e, r*) such that higher value of *g* corresponds to a higher likelihood that *R* binds to *E*. As mentioned above, we jointly learn separate epitope encoder *f*_*θ,e*_ and receptor encoder *f*_*θ,r*_ to compute *e* and *r*, respectively. We then utilize a contrastive learning objective to maximize the cosine similarity between the embeddings of epitopes and their receptors known to bind (Equation 2).

The loss function in ImmuneCLIP differs subtly from a conventional CLIP-style loss. Due to the nature of most binding assays for TCRs and pMHC complexes, there are more unique receptor sequences studied than there are unique epitope sequences, and there is often a many-to-one relationship between epitopes and receptors. Thus, in any given training minibatch of (epitope, TCR) pairs, a given epitope may occur more than once despite the receptors being unique. Rather than naively treating all “off-diagonal” epitope/TCR mismatches as negative samples, we construct target labels and utilize a binary cross entropy loss such that the embeddings are optimized across all known epitope/TCR pairs (Equation 3). The computed gradients are then back-propagated through the model, updating only the trainable LoRA matrices and projection layers.

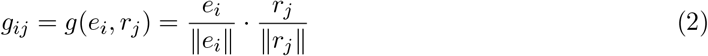

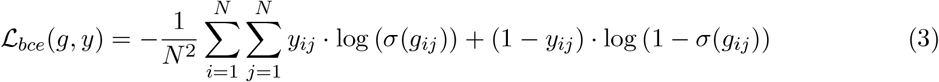

Additionally, in this specific setting where multiple receptors can bind to the same epitope sequence, batch size becomes an important hyperparameter in CLIP-style learning. A batch size that is too high may cause each batch to have redundant epitopes with high probability. A batch size that is too small may have a less diverse contrastive relationships among epitopes/receptors and can also slow training. Through empirical testing, a batch size of 16 was found to be ideal for training efficiency and downstream performance. Table S1 summarizes the hyperparameters tested for ImmuneCLIP as well as the optimal set of parameters used for model training.

## 3 Results

### 3.1 ImmuneCLIP Aligns TCR and Epitope Representations in a Biologically Meaningful Embedding Space

As mentioned in the previous section, ImmuneCLIP was trained on labeled pairs of epitope and TCR CDR3 *α, β* chain sequences. To evaluate ImmuneCLIP, we first investigated if the finetuned epitope and T-cell receptor encoders were successfully aligned in a biologically meaningful space. Specifically, we used ImmuneCLIP to screen the receptors in our test set and evaluated the model’s ability to recover each receptor’s known binding epitope. ImmuneCLIP was trained with the objective of maximizing the cosine similarity between receptor and TCR embeddings that are likely to bind in the human immune system. Thus, given an unseen receptor sequence and a pool of epitope sequences, the cosine similarity between the receptor’s embedding and an epitope’s embedding should correlate with their likelihood of binding *in vivo*. For each receptor, we computed the pairwise cosine similarity of its receptor with all epitopes, ranked the epitopes by their computed scores, and evaluated if the receptor’s paired epitope was in the top-*k* ranks at various thresholds (1, 5, 10).

Generally, immune receptors with similar CDR3 sequences tend to have similar epitope binding profiles [38, 39, 40, 41]. Thus, to ensure that ImmuneCLIP learned more biologically meaningful information than sequence similarity, we compared our model to Levenshtein distance (a baseline sequence similarity metric) and tcrdist3 (a state-of-the-art sequence alignment-based distance metric). Unlike our model, Levenshtein and tcrdist3 compute distances between TCR sequences alone and cannot directly compare receptors to epitopes. As a result, for every unseen receptor sequence in the test set, we compute its distance to all seen receptors in the training set, rank the receptors by the inverse of their sequence distance, and then predict the epitopes associated with the top-*k* receptors.

As shown in Figure 3a, ImmuneCLIP achieves the best performance for ranking the correct epitope within the top-*k* ranks for each receptor in our test set. This improved performance was shown to hold across multiple thresholds of *k*. Overall, our method was able to outperform both baseline and state-of-the-art sequence-based distance metrics, implying that ImmuneCLIP implicitly learned more meaningful information for TCR/epitope interactions from the contrastive fine-tuning process.

**Figure 3:**
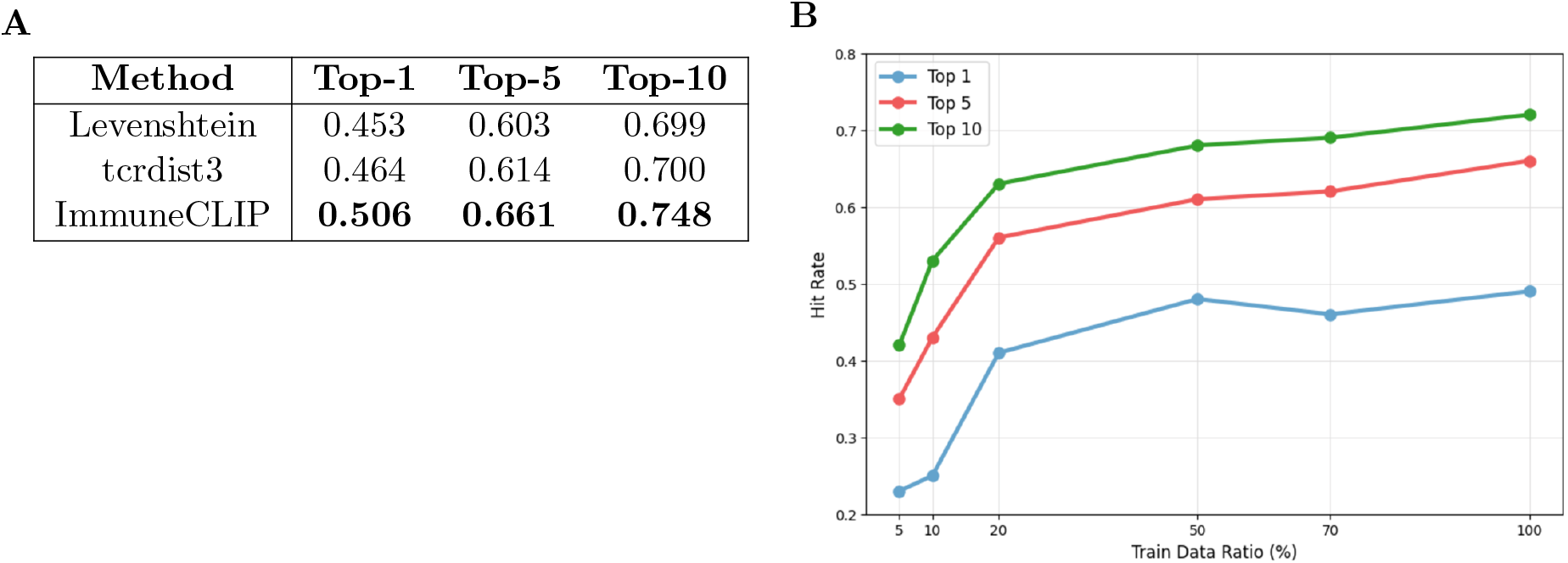
(a) Hit ratios of correct epitopes to test set TCRs for different methods at varying thresholds (*k* = 1, 5, 10), where we expect higher hit ratios as we increase the threshold. (b) Hit rate of ImmuneCLIP across varying training data fractions (5, 10, 20, 50, 70, and 100%) for Top-1, 5, 10 epitope ranking thresholds.

### 3.2 ImmuneCLIP is Effective in Binary Classification of Epitope/TCR Binding

Next, we evaluated ImmuneCLIP’s performance on the task of binary interaction prediction between unseen pairs of epitopes and T-cell receptors. For this task, instead of simply recovering the epitope for a TCR that is a known binder, a model must predict if a TCR will bind to a specific epitope or not. Binary binding labels can be extracted from ImmuneCLIP by treating the cosine similarity between a query receptor and epitope’s embeddings as logits. We benchmarked our model against TULIP [11] and STAPLER [10], both of which are deep learning methods that leverage sequence information from pMHC epitopes and TCR *α, β* chain CDR3 regions to predict binding labels. As a baseline, we also compared our method against sequence similarity-based methods Levenshtein and tcrdist3. Similar to Section 3.1, since we cannot directly compare epitopes and T-cell receptors with sequence distance, we executed the following method for prediction with Levenshtein distance and tcrdist3: for an unseen TCR *α, β* chain CDR3 and query epitope pair in the test set, we compute the distance from the unseen TCR to every seen TCR *α, β* chain CDR3 in our training set *that is known to bind to the query epitope* and find the smallest distance. The predicted binding score is then the inverse of the minimum distance.

We evaluated the performance of our model on the specific test sets curated for each of our comparison models (TULIP, STAPLER) in their papers. To prevent data leakage, we removed all epitope/TCR pairs in each of the test sets that our model had seen during training. To ensure a fair evaluation, we also excluded examples for any epitopes that had been seen by the other models during training but were not in our model’s training set, and vice versa. Lastly, a negligible number of test set epitopes could not be evaluated using Levenshtein and tcrdist3 because they had no corresponding training set examples, and thus we removed them as well. We found that ImmuneCLIP matched or exceed all other methods on both datasets in terms of AUROC and AUPR (Figure 4). In both test sets, we observe that the ROC curve for ImmuneCLIP bows more towards the left of the plot compared to others, indicating higher sensitivity and lower false positive rates.

**Figure 4:**
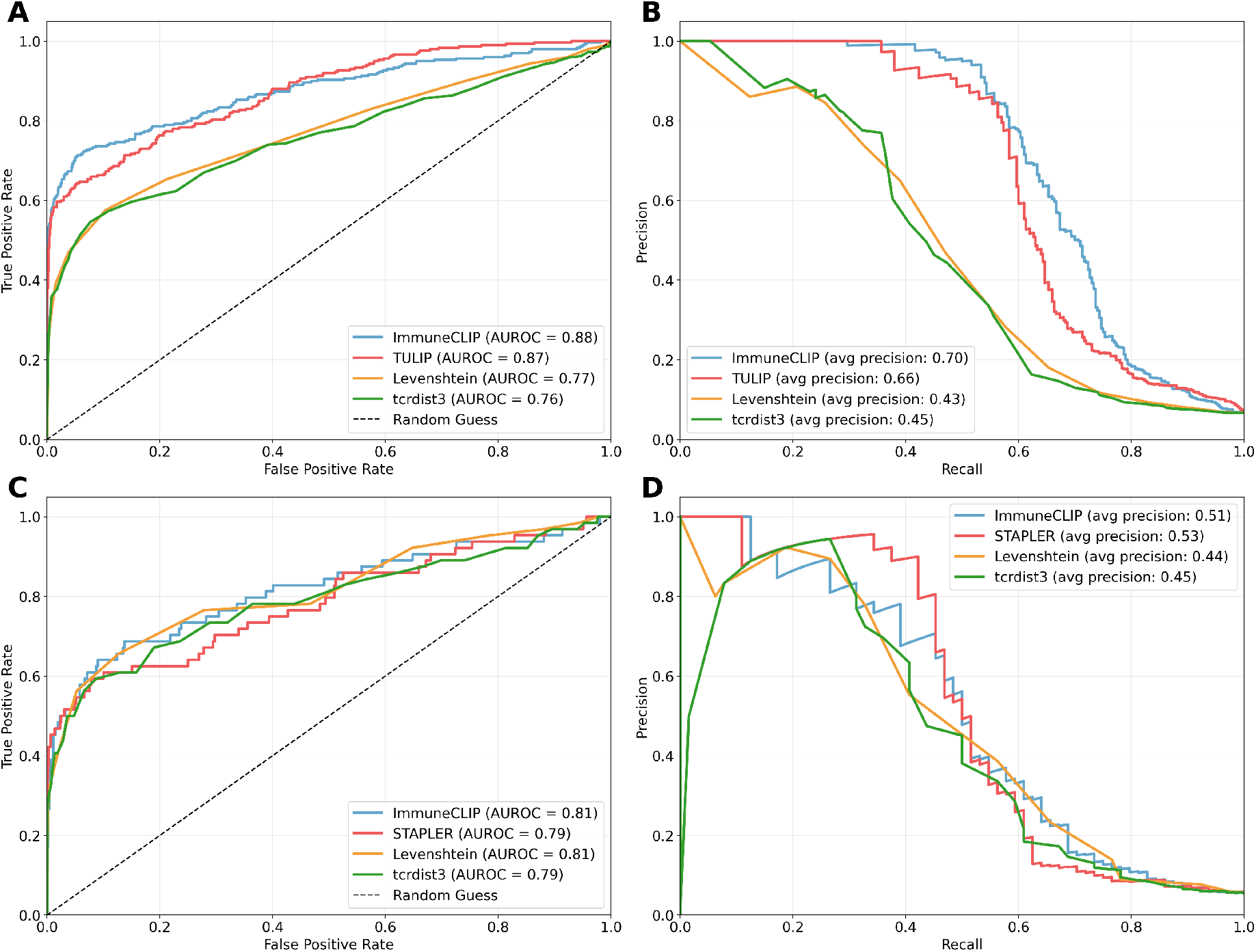
Performance comparison of ImmuneCLIP against TULIP (A, B) and STAPLER (C, D) on the epitope/TCR binding prediction task on the test sets from VDJdb as specified in Meynard-Piganeau et al. and Kwee et al., respectively.

### 3.3 Generalization to New Receptors from Few Shot

We sought to characterize the generalization capabilities of our model in the epitope ranking problem mentioned above. To do so, we conducted a series of experiments to measure test set model performance using varying amounts of training data. Our dataset consists of 146 unique epitopes, each associated with a set of known binding T-cell receptors, for a total of 8439 unique epitope/TCR pairs. As mentioned before, the pairs were partitioned into a training set (70%), validation set (15%) and a test set (15%). We created further subsets of the training data by sampling a fixed fraction of each epitope’s associated TCRs and trained models on the subsets to observe how ImmuneCLIP’s performance would scale with the amount of training data. Performance was evaluated on the same, unmodified test set in terms of hit rates across different top-*k* thresholds.

As illustrated in Figure 3b, the model’s hit rate improves as the included fraction of training data increases, particularly for the top-10 ranking. Notably, even when only 5% of the training data is included, ImmuneCLIP demonstrates a non-trivial ability to generalize, achieving hit rates of approximately 0.22, 0.30, and 0.41 for the top-1, top-5, and top-10 rankings, respectively. As the included fraction of training data increases to 100%, the model’s performance plateaus, indicating diminishing returns on hit rate improvement with additional data. These results highlight ImmuneCLIP’s capacity to generalize from limited data, suggesting its potential effectiveness in scenarios where only a small number of binding interactions are known. This is particularly promising for applications involving low-N epitope/TCR matching, where data scarcity is a common challenge.

### 3.4 Ablation Study

To access the effectiveness of various design choices within ImmuneCLIP, we conducted a comprehensive ablation study. The ImmuneCLIP framework consists of three main components: the pre-trained protein language model encoder, the fine-tuning methodology, and the projection layers. We examined each of these elements in detail.

We first explored the impact of the base PLM encoder on learning by training three variants of ImmuneCLIP, each with a different pre-trained LM as the TCR *α, β* CDR3 encoder. We tested ESM-2, a general protein language model trained on the UniRef protein sequence database, as well as TCR-BERT [18] and TCRLang-Paired [19], two specialized protein language models pretrained on human TRA and TRB sequences drawn from the public VDJdb [28] and PIRD [42] datasets and the OTS [19] dataset, respectively. As a control, we also replaced the TCR CDR3 encoder with a one-hot encoding followed by a simple MLP. For all experiments, we used the partial masking of tokens as described in Section 2.2 to minimize overfitting. We also up-sampled training examples with lower epitope counts to increase the model’s performance on low-N epitopes in the test set. The control one-hot encoding model yielded almost no learning, indicating that PLM pretraining is critical for performance. The application-specific foundation model TCR-BERT yielded the best downstream performance, followed by the general purpose protein language model ESM-2. Notably, TCRLang-Paired did not perform as well as TCR-BERT despite being equally specialized; we hypothesize that this is because TCRLang-Paired was pretrained on pairs of *full length* TCR *α, β* chains, while TCR-BERT was pretrained on individual TCR *α, β* CDR3 regions only. Since ImmuneCLIP fine-tunes PLMs using the CDR3 regions of TCRs, it is likely more in distribution with TCR-BERT.

**Table 1:** Performance comparison of different protein language models, fine-tuning strategies, and projection depths for ImmuneCLIP on the epitope ranking task. The table reports Top-1, 5, and 10 ranking accuracies, showing the impact of using pre-trained models, fine-tuning methods, and varying number of projection layers. The FT experiments were all conducted with 2-layer MLP projection on ESM-2 and TCR-BERT.

| Model Variant | Top-1 | Top-5 | Top-10 |
| --- | --- | --- | --- |
| **Embeddings** |  |  |  |
| One-hot Encoding | 0.002 | 0.052 | 0.080 |
| ESM-2 | 0.459 | 0.628 | 0.707 |
| TCR-BERT | **0.506** | **0.661** | **0.748** |
| TCRLang-Paired | 0.392 | 0.567 | 0.661 |
| **Fine-Tuning (FT)** |  |  |  |
| no FT (1.4M params) | 0.460 | 0.610 | 0.681 |
| Regular FT (768M params) | 0.499 | 0.650 | 0.715 |
| LoRA FT (2M params) | 0.511 | 0.653 | 0.718 |
| **Projection (on ESM-2)** |  |  |  |
| no projection (FT only) | 0.381 | 0.578 | 0.673 |
| 1 MLP layer | 0.459 | 0.628 | 0.707 |
| 2 MLP layers | 0.468 | 0.656 | 0.736 |

We also conducted ablations on different fine-tuning methods and projection depths. As a control, we found that completely freezing the PLM encoders and performing no fine-tuning still resulted in decent downstream performance, solely through updating the projection layer weights. However, with fine-tuning, we were able to achieve higher performance across all 3 thresholds in the epitope ranking task. Low rank adaptation achieved equivalent performance to full fine-tuning while dramatically reducing GPU memory usage (< 4 GB) and training time, demonstrating the value of parameter efficient fine-tuning [32]. We noticed a similar trend from the projection depth ablations; removing the projection layers and solely fine-tuning the PLM encoders yielded significantly lower performance. Overall, our study was able to highlight that both LoRA fine-tuning and projection contributed to the overall performance of ImmuneCLIP.

## 4 Discussion and Conclusion

In this study, we introduced ImmuneCLIP, a scalable approach for contrastively fine-tuning protein language models (PLMs) to learn epitope/TCR binding interactions across multiple epitopes. By embedding and aligning both TCR and epitope sequences in a shared latent space, our method offers a purely sequence-based solution to the challenge of predicting multi-epitope TCR/pMHC interactions. We demonstrate that ImmuneCLIP outperforms traditional sequence similarity metrics and state-of-the-art deep learning models like TULIP and STAPLER while capturing biologically meaningful features. In particular, ImmuneCLIP shows superior performance to Levenshtein distance and tcrdist3 on epitope ranking, indicating that the model has learned critical features related to binding interactions beyond simple measures of similarity. Additionally, ImmuneCLIP ‘s strong results on the epitope/receptor binding prediction task supports the value of contrastive fine-tuning for accurately identifying TCR/epitope pairs. The model’s ability to generalize well, even with limited training data, is particularly promising for applications where data scarcity is a limiting factor, such as rare epitopes or low-N epitope/TCR matching.

Despite these promising results, several limitations remain. Although the dataset used in this study is large, it includes a limited variety of unique epitopes, which may restrict the model’s generalizability to rare or highly diverse epitopes. Expanding the epitope dataset would likely improve both performance and robustness. Additionally, while ImmuneCLIP performs well with sequence-based embeddings, integrating structural data—such as the 3D conformations of TCR/pMHC complexes—could enhance predictive accuracy, which we plan to explore in future work. The model’s performance plateau with the current dataset suggests that advanced pre-training techniques or data augmentation strategies could further improve results, particularly when additional data collection is not feasible. Finally, although ImmuneCLIP demonstrates some ability to generalize based on epitope similarity, it struggles with unseen, novel epitopes. As shown in Figure S1, the model performs better on test epitopes that share sequence similarity with those in the training set, underscoring the need for further improvements to handle truly novel epitopes.

Looking ahead, ImmuneCLIP presents several exciting opportunities for future research and applications. Its flexibility and performance on sequence-based data suggest that it could be adapted for use with other immune receptor families which face similar challenges in predicting binding interactions given sparse structural data. Additionally, expanding ImmuneCLIP to predict quantitative affinity, rather than binary binding outcomes, could offer more nuanced insights for therapeutic applications, including drug discovery and optimization. With growing immune receptor datasets, ImmuneCLIP has the potential to map extensive receptor-target interaction networks, provide deeper insights into immune system function, and enable more targeted therapeutic interventions.

## Code and Data Availability

The dataset for training and evaluating ImmuneCLIP was downloaded from the public repository of MixTCRPred (https://github.com/GfellerLab/MixTCRpred). The source code for training and inference are available at https://github.com/kundajelab/ImmuneCLIP.

## Appendix

**Figure S1:**
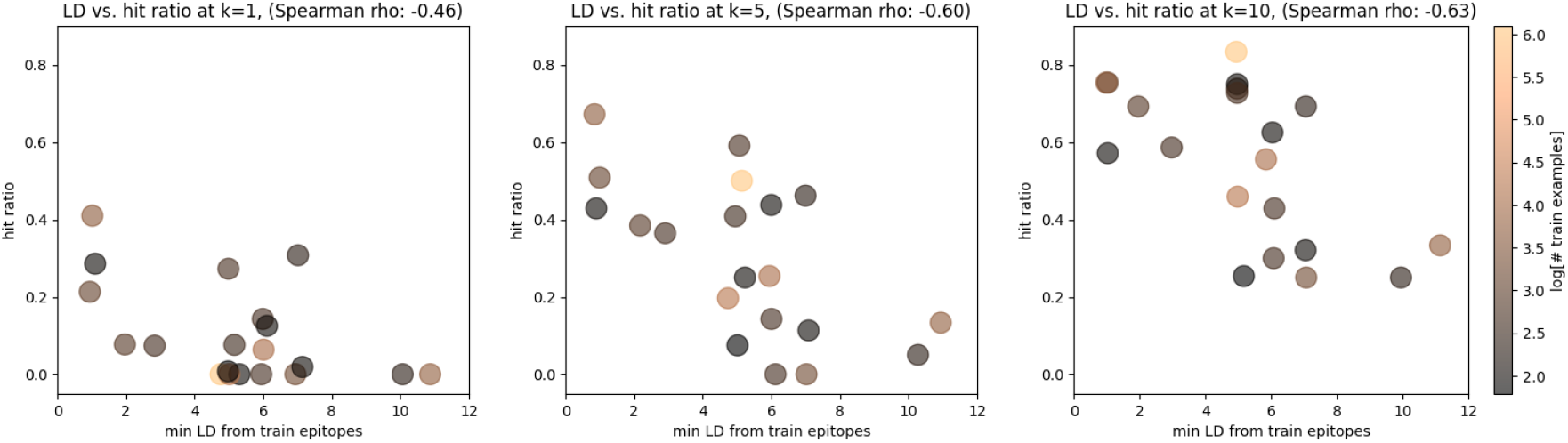
Generalization performance of ImmuneCLIP on unseen epitopes across different ranking thresholds (*k* = 1, 5, 10). For each unseen epitope in test set, we plotted the minimum Levenshtein distance (LD) to epitopes in the training set against the hit accuracy, where we would *expect* the accuracy (hit ratio) to drop as sequences become more dissimilar. The model performs better when test epitopes share sequence similarity with the training set, but struggles with entirely novel epitopes, highlighting the challenge of full generalization.

**Table S1:** Hyperparameter search space for ImmuneCLIP, including variations in masking, upsampling, LoRA rank, learning rates, batch sizes, non-linearities, PLMs in the TCR encoder, and projection layer configurations. The optimal choice of parameters chosen for model training is in **bold**.

| Parameter | Search Space |
| --- | --- |
| Using making of sequence during training | **Yes**, No |
| Using upsampling of low-N epitopes | **Yes**, No |
| LoRA Rank | 4, **8** |
| Learning rates | 0.01, **0.001**, 0.0001, 0.00005 |
| Batch size | 8, **16**, 32 |
| Non-linearities | **LeakyReLU** |
| PLM in TCR Encoder | ESM-2, **TCR-BERT**, TCRLang |
| Projection layers | 1, **2** |
| Projection dimension | 256, **512**, 1024 |

